# Modeling Sepsis-Associated ARDS Using a Lung Endothelial Microphysiological System

**DOI:** 10.1101/2023.10.10.561102

**Authors:** Nai-Wen Liang, Carole Wilson, Brooke Davis, Ian Wolf, Tonela Qyli, Joy Moy, David J. Beebe, Lynn M. Schnapp, Sheena C. Kerr, Hilary E. Faust

**Author notes:** Funding: Dr. Faust is funded by KL2TR002374 and UL1TR002373 awarded to UW ICTR through NIH NCATS. Competing interests: David J. Beebe holds equity in Bellbrook Labs LLC, Tasso Inc., Salus Discovery LLC, Lynx Biosciences Inc., Stacks to the Future LLC, Flambeau Diagnostics LLC, and Onexio Biosystems LLC.

## Abstract

Acute respiratory distress syndrome due to non-pulmonary causes exhibits prominent endothelial activation which is challenging to assess in critically ill patients. Preclinical *in vivo* models of sepsis and ARDS have failed to yield useful therapies in humans, perhaps due to interspecies differences in inflammatory responses. Use of microphysiological systems (MPS) offer improved fidelity to human biological responses and better predict pharmacological responses than traditional culture. We adapted a lung endothelial MPS based on the LumeNEXT platform to evaluate the effect of plasma from critically ill sepsis patients on endothelial permeability, adhesion molecule expression and inflammatory cytokine production. Lumens incubated with sepsis plasma exhibited areas of contraction, loss of cellular coverage, and luminal defects. Sepsis plasma-incubated lumens had significantly increased permeability compared to lumens incubated with healthy donor plasma. ICAM-1 expression increased significantly in lumens incubated with sepsis plasma compared with those incubated with healthy control plasma, while concentrations of IL-6, IL-18, and soluble VEGF-R1 increased in sepsis plasma before and after incubation in the MPS compared with healthy control plasma. Use of the lung endothelial MPS may enable interrogation of specific mechanisms of endothelial dysfunction that promote ARDS in sepsis patients.

---

Acute respiratory distress syndrome (ARDS) complicates 10% of sepsis cases and continues to have unacceptably high mortality without any available targeted therapies.^1, 2^ Mechanisms contributing to sepsis-associated ARDS prominently feature endothelial dysfunction resulting in loss of barrier function, extravasation of immune cells, and microthrombosis.^3^ Identifying specific targetable mediators of endothelial dysfunction is a critical yet elusive objective in achieving precision medicine approaches to prevention and treatment of ARDS.

Preclinical *in vivo* models of sepsis and ARDS have failed to yield useful therapies in humans, perhaps due to interspecies differences in inflammatory responses.^4, 5^ Efforts to understand human endothelial responses to sepsis contributing to ARDS are constrained, however, by limited access to endothelial tissue in critically ill patients and a lack of *in vitro* models that convincingly mimic lung vasculature.^6^ New approaches are needed to understand mechanisms of endothelial activation in sepsis in order to identify and evaluate targeted therapies prior to clinical trials in humans.

Three-dimensional lung endothelial microphysiological systems (MPS), sometimes referred to as a “lung on a chip,” may fill this gap by offering insight into mechanisms of endothelial dysfunction and responses to therapies. Culture of endothelial cells in a physiologically relevant geometry can have a profound impact on cellular function. MPS have been shown to offer improved fidelity to human biological responses and better predict pharmacological responses than traditional culture, yielding significant advantages for evaluating human therapeutics.^7, 8^ We adapted an existing lung endothelial MPS, based on the LumeNEXT platform, to evaluate lung microvascular endothelial responses to plasma from sepsis patients (Supplemental Figure 1).^9^ We specifically interrogated the effect of plasma from critically ill sepsis patients on endothelial barrier function, inflammatory cytokine production and vascular activation markers.

The study was approved by the University of Wisconsin Institutional Review Board (IRB 2021-0974). We obtained plasma from ambulatory control subjects (“normal”) and from patients enrolled in the University of Wisconsin Sepsis Biobank, utilizing residual clinical lab specimens from the sample most proximal to ICU admission. Written informed consent was obtained from patients or their surrogates. Patient characteristics are summarized in Table 1. Plasma was diluted 1:5 and 1.6 ug/mL heparin was added to prevent coagulation. LumeNEXT devices were fabricated as previously described^9, 10^ and filled with a collagen/fibronectin hydrogel to mold lumens which were seeded with lung microvascular endothelial cells (Lonza, Basal, Switzerland). Devices cultured in endothelial growth medium (EGM)-2 MV culture medium (Lonza) until confluence. Diluted plasma was incubated in the endothelial lumens for 16 hours and then collected. Lumen barrier function was assessed by measuring diffusion of Texas Red-labeled dextran from the vascular lumen. Permeability was quantified as the area under the curve (AUC) of the change in fluorescence from 0 to 10 minutes of dextran incubation normalized to the minimum (0) and maximum (1) of fluorescence at time 0. Lumens were stained with nuclear (Hoechst, Thermo Fisher Scientific, Waltham, MA) and anti-F actin (CellMask, Thermo Fisher) and imaged using a Nikon TI Eclipse microscope to assess lumen integrity and cellular confluence. Lumens were also stained with intracellular adhesion molecule 1 (ICAM-1, R&D Systems, Minneapolis, MN) to determine the effect of sepsis plasma on vascular adhesion markers. We quantified cytokine and endothelial activation markers on plasma before and after incubation in the MPS lumens using a custom multiplexed bead-based ELISA (Luminex, Thermo Fisher).

**Table 1.**

| Demographic characteristics (n=10) |  |
| --- | --- |
| Race |  |
| Black | 0 (0%) |
| White | 10 (100%) |
| Asian | 0 (0%) |
| Other | 0 (0%) |
| Age, y | 64 (49-80)* |
| Female sex | 2 (20%) |
| Male sex | 8 (80%) |
| Illness and injury severity |  |
| Pulmonary source | 2 (20%) |
| Pre-ICU shock | 3 (30%) |
| Medical History |  |
| Diabetes | 2 (20%) |
| HTN | 6 (60%) |
| CHF | 3 (30%) |
| CKD | 3 (30%) |
| Outcomes |  |
| ARDS | 3 (30%) |
| 90-day mortality | 4 (40%) |
| * Interquartile range |  |

Incubation of sepsis plasma induced marked cellular changes in endothelial lumens compared with incubation of healthy donor plasma (Figure 1A). Lumens incubated with sepsis plasma exhibited areas of contraction, loss of cellular coverage, and luminal defects. Sepsis plasma-incubated lumens had significantly increased permeability (area under the curve [AUC] 477.31, interquartile range [IQR] 63.33) compared to lumens incubated with healthy donor plasma (363.10, IQR 14.88, p=0.000027, Figure 1B). ICAM-1 expression increased significantly in lumens incubated with sepsis plasma compared with those incubated with healthy control plasma (0.684, IQR 0.15 vs 1.042, IQR 0.071, p=0.0000032, Figure 1C and 1D). Concentrations of interleukin (IL)-6, IL-18, and soluble vascular endothelial growth factor-receptor 1 (VEGF-R1) increased in sepsis plasma before and after incubation in the MPS compared with healthy control plasma that did not reach statistical significance (Figure 1E, Supplemental Table 1). Changes in pre-and post-incubation concentrations of additional cytokines and vascular adhesion markers are shown in Supplemental Figure 2 and Supplemental Table 1.

**Figure 1.**
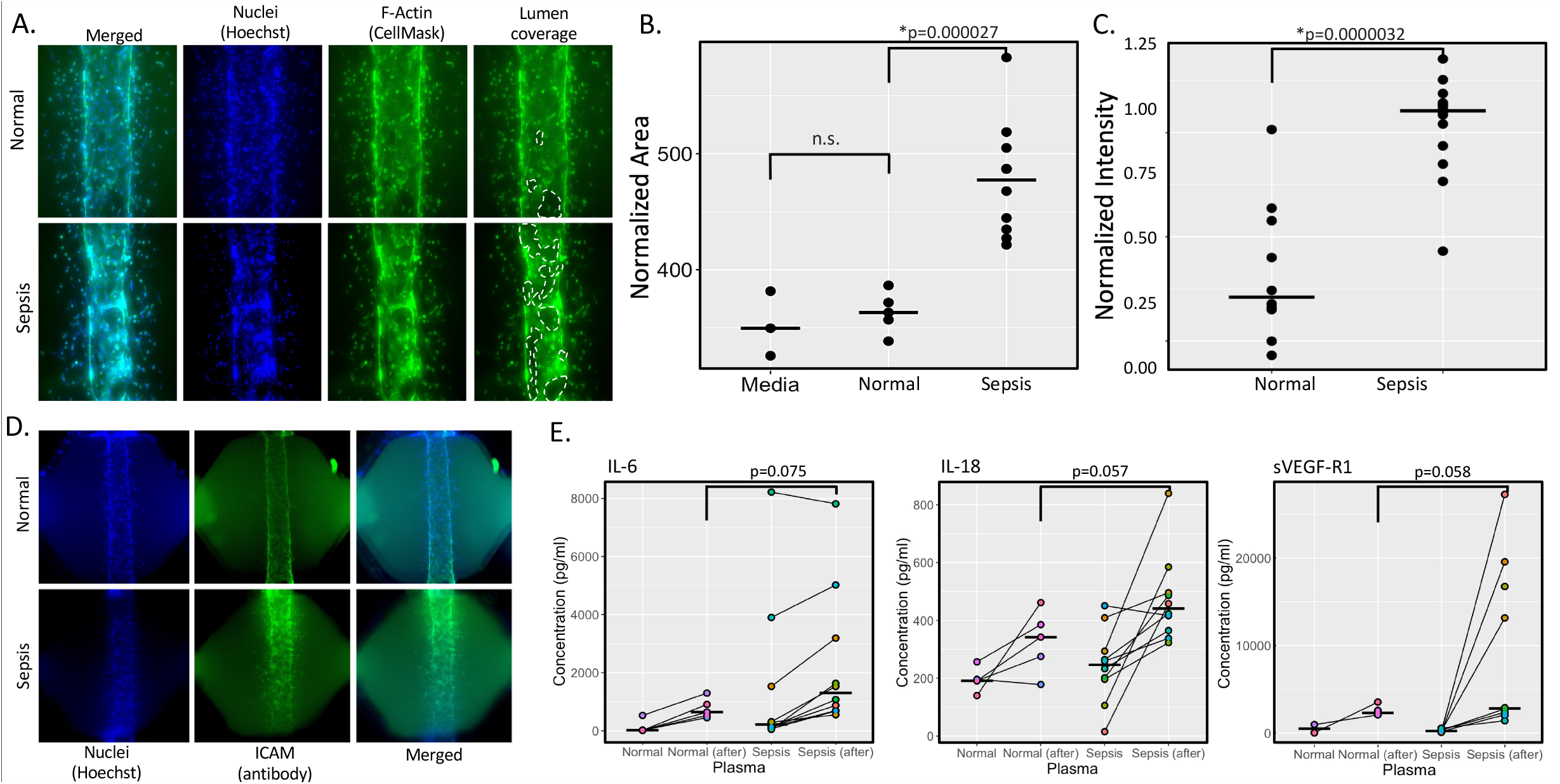
Sepsis Plasma Induces Endothelial Dysfunction in a Lung Endothelial MPS Lung endothelial MPS incubated with healthy donor plasma (“normal”) and sepsis patient plasma (“sepsis) for 16 hours. A) Staining of lumens incubated with normal and sepsis plasma. B) Increased vascular permeability in lumens incubated with sepsis plasma (n=10 sepsis patients) compared with normal plasma (n=5 healthy controls). C) Intensity of ICAM staining after incubation with representative normal (n=10 technical replicates) and sepsis plasma (n=15 technical replicates) indicating significantly increased ICAM expression on sepsis lumens. D) ICAM staining on representative sepsis and normal lumens. E) Inflammatory and vascular activation cytokines were numerically but not statistically increased in residual sepsis plasma compared with healthy plasma after 16 hours of incubation.

In conclusion, we present the first known use of sepsis plasma in a lung endothelial MPS model of sepsis-associated ARDS. Lung endothelial dysfunction is thought to be critical to the development of ARDS triggered by circulating factors released during systemic inflammatory states such as sepsis, but specific mechanisms active in humans are challenging to study in critically ill patients. We demonstrated morphologic, functional, and secreted alterations in endothelial cells exposed to sepsis plasma compared with healthy donor plasma. We reproduced barrier dysfunction and upregulation of adhesion molecules, while also showing numerical increases in cytokines associated with activation of specific inflammatory pathways. Increases in ICAM-1 expression are consistent with the induction of endothelial activation, while the trend towards increased IL-6 before and after incubation are similar to molecular profiles demonstrated in human studies of extrapulmonary ARDS.^3^ By replicating these processes in an accessible *in vitro* environment utilizing human endothelial cells, we can directly assess causality of specific candidate mechanisms and establish the utility of targeted therapies *in vitro*. For example, the trend towards increased IL-18 induced by sepsis plasma suggests activation of inflammasome signaling, which could motivate targeted inhibition for prevention or treatment of sepsis-associated ARDS. This approach may represent a rational strategy to screen potential preventive and therapeutic agents safely prior to undertaking costly and time-consuming clinical trials in critically ill ARDS patients.

