## Supplemental Data for "Modeling Sepsis-Associated ARDS Using a Lung Endothelial Microphysiological System"

### Slide 1
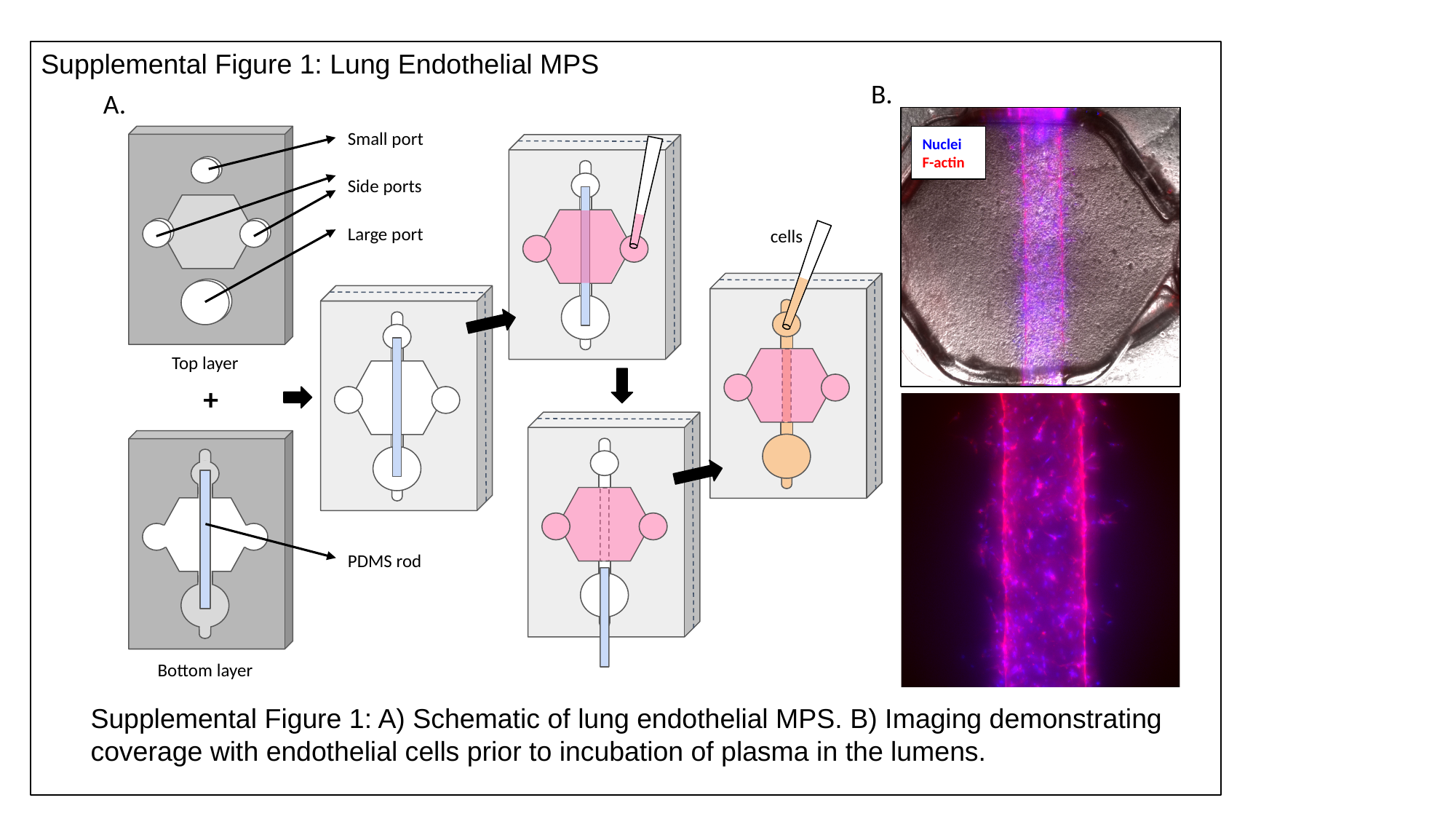

Supplemental Figure 1: Lung Endothelial MPS
B.
A.
Small port
Nuclei
F-actin
Side ports
Large port
cells
Top layer
+
PDMS rod
Bottom layer
Supplemental Figure 1: A) Schematic of lung endothelial MPS. B) Imaging demonstrating coverage with endothelial cells prior to incubation of plasma in the lumens.

### Slide 2
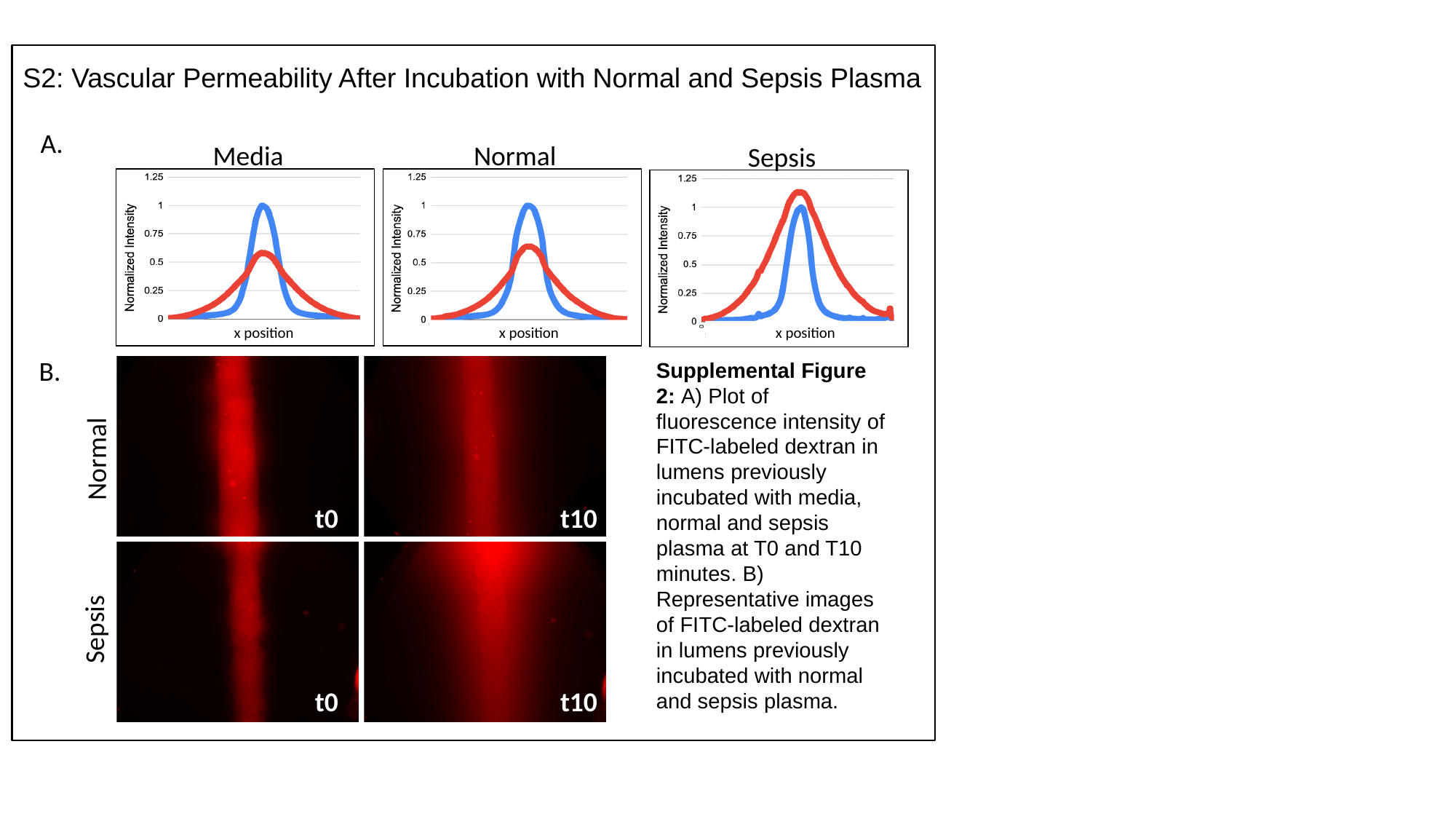

#### S2: Vascular Permeability After Incubation with Normal and Sepsis Plasma
A.
Media
Normal
Sepsis
x position
x position
x position
B.
Supplemental Figure 2: A) Plot of fluorescence intensity of FITC-labeled dextran in lumens previously incubated with media, normal and sepsis plasma at T0 and T10 minutes. B) Representative images of FITC-labeled dextran in lumens previously incubated with normal and sepsis plasma.
Normal
t0
t10
Sepsis
t0
t10

### Slide 3
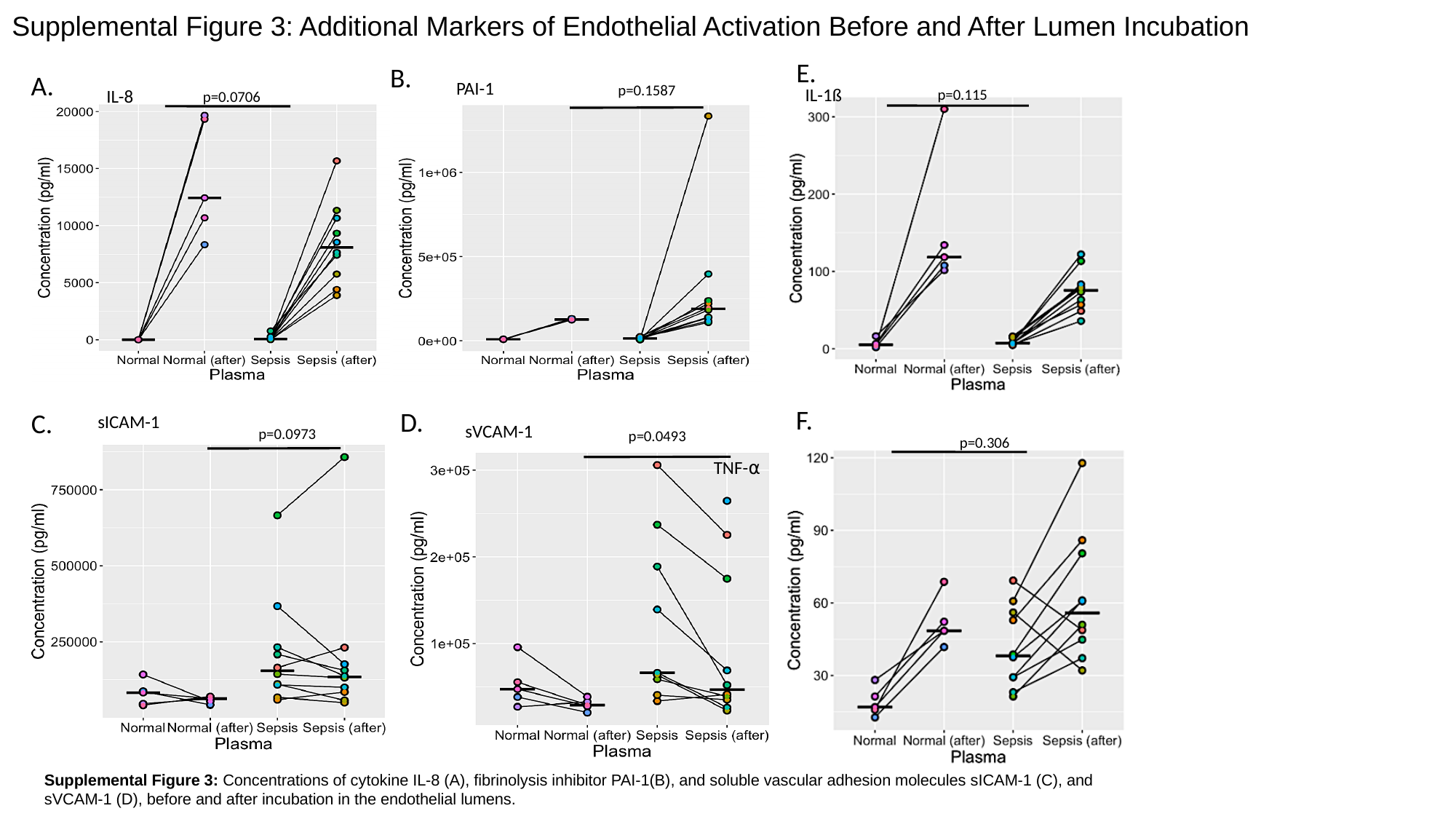

#### Supplemental Figure 3: Additional Markers of Endothelial Activation Before and After Lumen Incubation
E.
B.
A.
PAI-1
p=0.1587
IL-8
IL-1ß
p=0.115
p=0.0706
F.
sICAM-1
p=0.0973
D.
C.
sVCAM-1
p=0.0493
p=0.306
TNF-⍺
Supplemental Figure 3: Concentrations of cytokine IL-8 (A), fibrinolysis inhibitor PAI-1(B), and soluble vascular adhesion molecules sICAM-1 (C), and sVCAM-1 (D), before and after incubation in the endothelial lumens.

### Slide 4
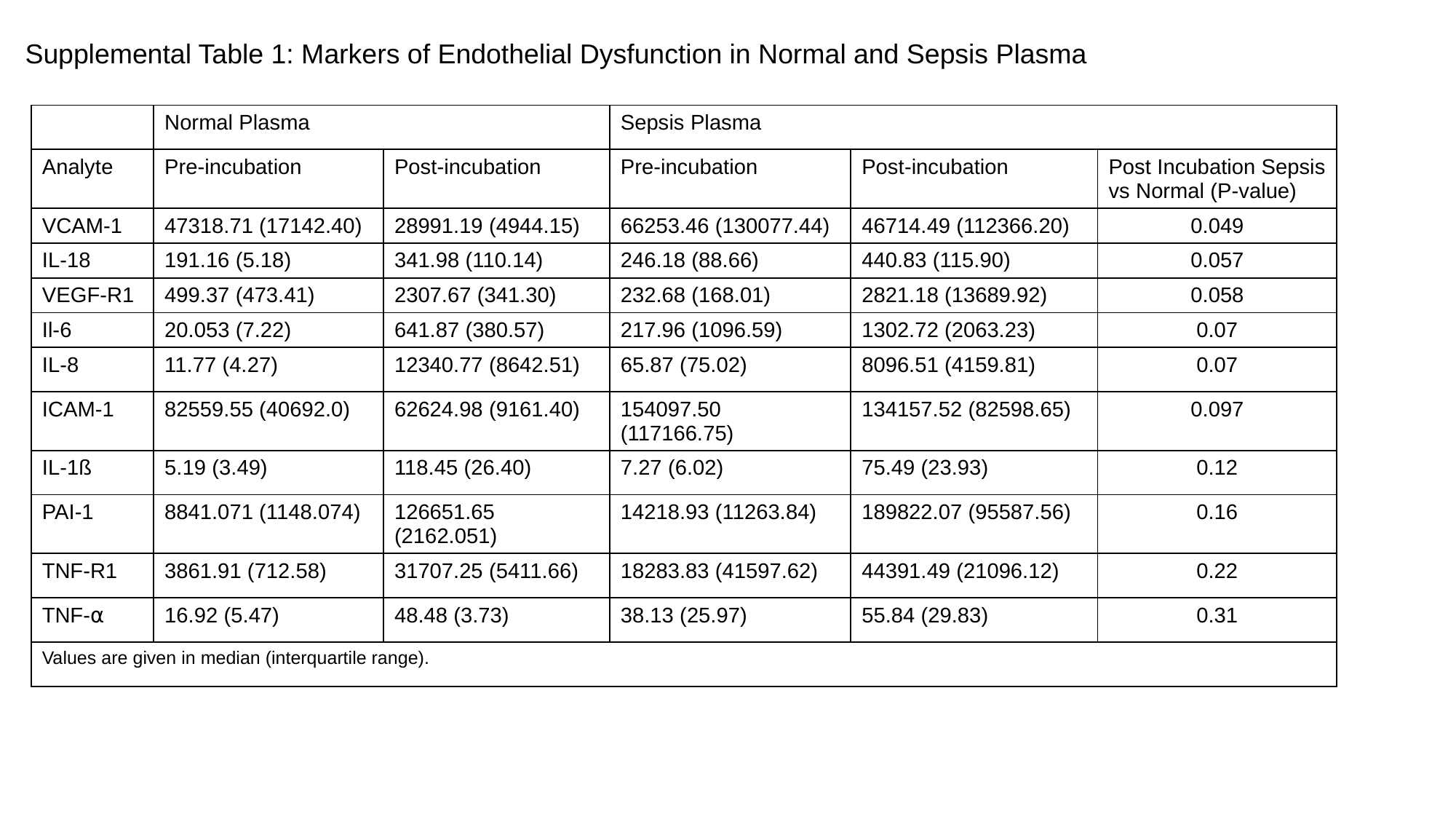

#### Supplemental Table 1: Markers of Endothelial Dysfunction in Normal and Sepsis Plasma
| | Normal Plasma | | Sepsis Plasma | | |
| --- | --- | --- | --- | --- | --- |
| Analyte | Pre-incubation | Post-incubation | Pre-incubation | Post-incubation | Post Incubation Sepsis vs Normal (P-value) |
| VCAM-1 | 47318.71 (17142.40) | 28991.19 (4944.15) | 66253.46 (130077.44) | 46714.49 (112366.20) | 0.049 |
| IL-18 | 191.16 (5.18) | 341.98 (110.14) | 246.18 (88.66) | 440.83 (115.90) | 0.057 |
| VEGF-R1 | 499.37 (473.41) | 2307.67 (341.30) | 232.68 (168.01) | 2821.18 (13689.92) | 0.058 |
| Il-6 | 20.053 (7.22) | 641.87 (380.57) | 217.96 (1096.59) | 1302.72 (2063.23) | 0.07 |
| IL-8 | 11.77 (4.27) | 12340.77 (8642.51) | 65.87 (75.02) | 8096.51 (4159.81) | 0.07 |
| ICAM-1 | 82559.55 (40692.0) | 62624.98 (9161.40) | 154097.50 (117166.75) | 134157.52 (82598.65) | 0.097 |
| IL-1ß | 5.19 (3.49) | 118.45 (26.40) | 7.27 (6.02) | 75.49 (23.93) | 0.12 |
| PAI-1 | 8841.071 (1148.074) | 126651.65 (2162.051) | 14218.93 (11263.84) | 189822.07 (95587.56) | 0.16 |
| TNF-R1 | 3861.91 (712.58) | 31707.25 (5411.66) | 18283.83 (41597.62) | 44391.49 (21096.12) | 0.22 |
| TNF-⍺ | 16.92 (5.47) | 48.48 (3.73) | 38.13 (25.97) | 55.84 (29.83) | 0.31 |
| Values are given in median (interquartile range). | | | | | |

### Slide 5
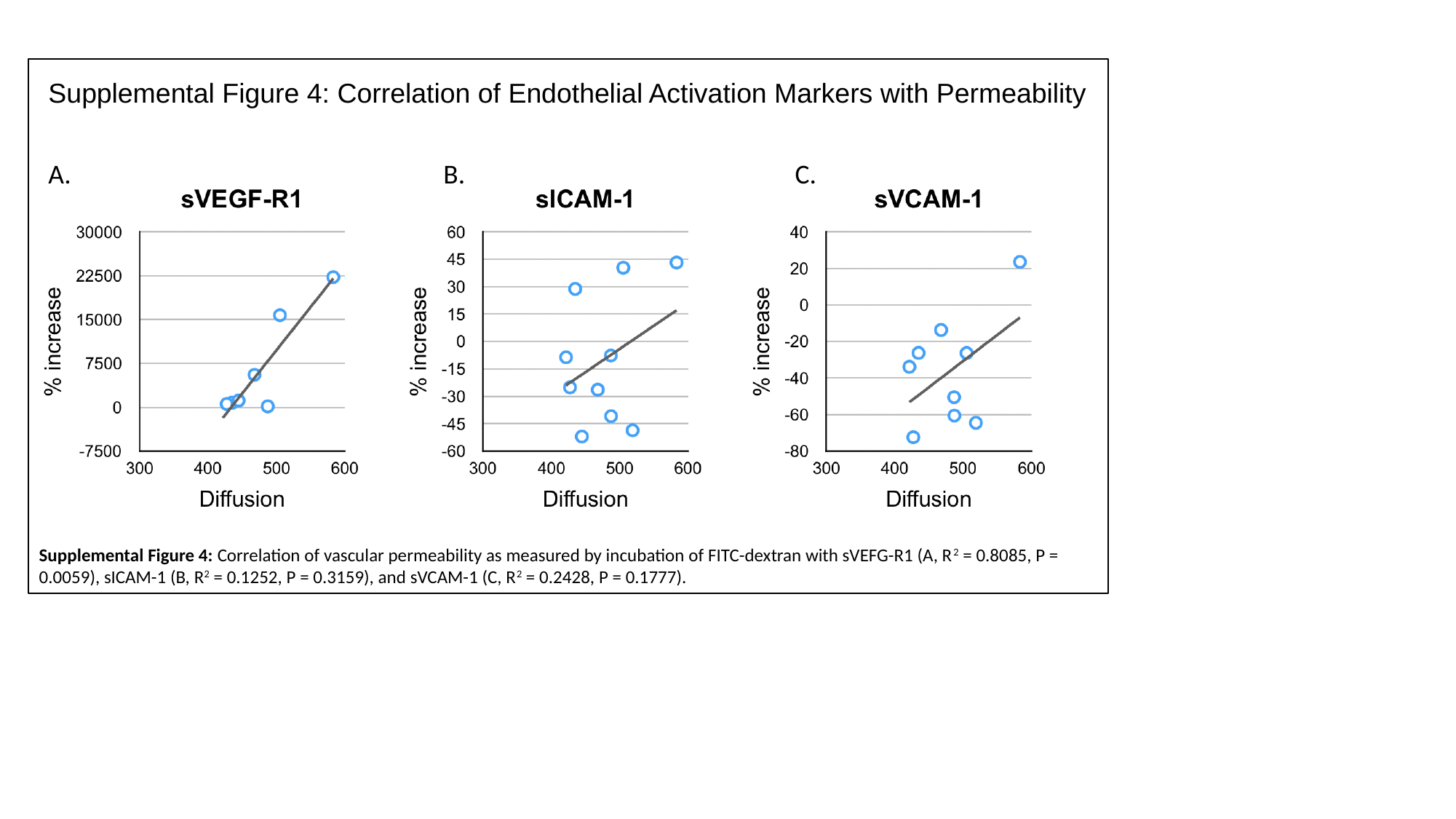

#### Supplemental Figure 4: Correlation of Endothelial Activation Markers with Permeability
B.
C.
A.
=
Supplemental Figure 4: Correlation of vascular permeability as measured by incubation of FITC-dextran with sVEFG-R1 (A, R^2^ = 0.8085, P = 0.0059), sICAM-1 (B, R^2^ = 0.1252, P = 0.3159), and sVCAM-1 (C, R^2^ = 0.2428, P = 0.1777).
